# Combined EGFR and ROCK inhibition in TNBC leads to cell death via impaired autophagic flux

**DOI:** 10.1101/661272

**Authors:** Stamatia Rontogianni, Sedef Iskit, Sander van Doorn, Daniel S. Peeper, A. F. Maarten Altelaar

**Affiliations:** Biomolecular Mass Spectrometry and Proteomics, Bijvoet Center for Biomolecular Research and Utrecht Institute for Pharmaceutical Sciences, Utrecht University, Padualaan 8, 3584 CH Utrecht, The Netherlands; Netherlands Proteomics Center, Padualaan 8, 3584 CH Utrecht, The Netherlands; Division of Molecular Oncology, The Netherlands Cancer Institute, 1066 CX Amsterdam, the Netherlands; Mass Spectrometry and Proteomics Facility, The Netherlands Cancer Institute, 1066 CX Amsterdam, the Netherlands

## Abstract

Triple-negative breast cancer (TNBC) is an aggressive subtype of breast cancer with very limited therapeutic options. We have recently shown that the combined inhibition of EGFR and ROCK in TNBC cells results in cell death, however, the underlying mechanisms remain unclear. To investigate this, here we applied a mass spectrometry-based proteomic approach to identify proteins altered upon single and combination treatments. Our proteomic data revealed autophagy as the major molecular mechanism implicated in the cells’ response to combinatorial treatment. In particular, we here show that EGFR inhibition by gefitinib treatment alone induces autophagy, a cellular recycling process that acts as a cytoprotective response for TNBC cells. However, combined inhibition of EGFR and ROCK leads to autophagy blockade and accumulation of autophagic vacuoles. Our data show impaired autophagosome clearance as a cause of antitumor activity. We propose that the inhibition of the autophagic flux upon combinatorial treatment is attributed to the major cytoskeletal changes induced upon ROCK inhibition, given the essential role the cytoskeleton plays throughout the various steps of the autophagy process.

## Introduction

Triple-negative breast cancer (TNBC), comprising 10-20% of all breast cancers, is an aggressive subtype of breast cancer, which is typically associated with poor prognosis. TNBC tumors are immunohistochemically defined by a lack of estrogen receptor (ER) and progesterone receptor (PR) expression as well as human epidermal growth factor receptor 2 (HER2) amplification and are therefore, insensitive to the established hormonal therapy and/or HER2 targeted treatment. Although it is possible to treat TNBC by surgery and systemic therapy, treatment-resistant recurrences are common [1]. Despite extensive research, the absence of hormone receptors or a common genetic vulnerability has prevented the development of a clinically established targeted treatment against TNBC [2]. Therefore, the development of new targeted therapies for patients with TNBC are urgently needed [3].

In TNBC, the epidermal growth factor receptor (EGFR) is frequently overexpressed, making it a potential therapeutic target. Currently, two types of EGFR inhibitors are being used in the clinic, small molecular tyrosine kinase inhibitors and monoclonal antibodies, which have already proven effective in other types of cancers, such as colorectal cancer. Unfortunately, no EGFR therapies are currently approved for TNBC because of low response rates, necessitating better markers for patient stratification [4] as well as the exploration of combination therapies [5, 6]. In this light, two independent studies recently showed great synergistic antitumor activity when inhibitors of the RAF-MEK-ERK cascade were combined with autophagy inhibition in pancreatic and other RAS-driven cancers [7, 8].

Macroautophagy (hereafter referred to as autophagy) is a highly dynamic multi-step biological process of self-cannibalization that involves the degradation of damaged organelles, misfolded proteins and long-lived macromolecules in lysosomes. This process occurs under basal conditions for example, to degrade long-lived proteins, but is drastically elevated in cells under stress, such as starvation or hypoxia, as a protective mechanism, allowing cells to survive [9]. Autophagy, which is characterized by the engulfment of cargo molecules by double-membrane vesicles, called autophagosomes, is an orchestrated process involving several steps. It starts with the formation and elongation of the phagophore, which enwraps and sequesters portions of the cytoplasm containing autophagic substrates, and then it expands through acquisition of lipids, and ultimately seals to generate a completed double membrane called autophagosome. Following closure, the autophagosome fuses with the lysosome to form the autolysosome, where the sequestered cargo is degraded and recycled [10, 11].

In the context of cancer, the activation of autophagy is considered to be a double-edged sword. On the one hand, it functions primarily as a tumor suppressor mechanism, by clearing damaged organelles, maintaining cell homeostasis and protecting normal cell growth [12]. Conversely, in established cancers, autophagy may become a key survival mechanism for tumor cells under a variety of stresses. For instance, evolving tumors develop regions of hypoxia and nutrient limitation, and under such harsh conditions, cancer cells adapt by inducing autophagy to protect themselves from cell death [13]. Moreover, a growing body of evidence suggests that autophagy activation plays a cytoprotective role in cancer cells undergoing various anti-cancer treatments, resulting in poor treatment outcomes and the development of treatment resistance [14]. Accordingly, preclinical studies have shown that genetic or pharmacological inhibition of cytoprotective autophagy can overcome therapy resistance and promote tumor regression [15-17].

We previously carried out *in vivo* and *in vitro* screens complemented with pharmacologic screens to identify drug combinations that effectively impair TNBC cell growth. We reported that combined inhibition of EGFR and ROCK induces cell cycle arrest in TNBC cells [18]. However, the underlying mechanisms by which co-inhibition of EGFR and ROCK induces TNBC cell death remain unclear. Here, we set out to elucidate the synergistic effect of the combined treatment using mass spectrometry-based quantitative (phospho)proteomics. We employed a two-dimensional proteomic strategy by combining offline high-pH reversed phase fractionation with nanoLC-MS/MS for deep proteomic profiling in order to identify proteins and pathways altered upon single and combination treatments. Interestingly, our data showed a significant increase in the expression levels of autophagy-related proteins upon EGFRi-treatment, both at the proteome and phosphoproteome level, while combined treatment with EGFRi and ROCKi leads to impaired autophagy, resulting in increased cell death.

## Results

### Proteomic profiling of TNBC cells upon single and combination treatments

To gain understanding of the mechanisms underlying the synergistic effect of the combined inhibition of EGFR and ROCK in triple-negative breast cancer cells we performed a mass spectrometry-based proteomics analysis. As a model system for our study, we selected the triple-negative breast cancer cell line Hs578T, to identify proteomic differences in signaling upon treatment with either of the two single inhibitors and their combination treatment. Thus, Hs578T cells were treated either with DMSO (control), gefitinib (EGFRi), GSK269962A (ROCKi) or with their combination treatment (EGFRi+ROCKi). Consistent with our previous findings [18], EGFRi+ROCKi inhibition significantly impaired the TNBC cell growth compared to the EGFRi and ROCKi alone (Figure 1A). To gain insight into the global signaling changes occurring across the different treatments, we employed a label-free quantitative (phospho)proteomics approach. Briefly, cells were treated for 48h, lysed and subsequently the protein extracts were in-solution digested by LysC/trypsin. To obtain a deep proteome coverage, we generated five fractions by off-line high-pH reverse phase chromatography (HpH) [19]. In parallel, for the phosphoproteomics analysis, an automated phosphopeptide enrichment step was performed using Fe(III)-IMAC cartridges on an AssayMAP Bravo platform as has been previously described [20]. All samples were analyzed by nanoLC-MS/MS coupled to a quadrupole Orbitrap high resolution mass spectrometer (Q Exactive Plus) followed by data analysis in MaxQuant. In total, we identified 7,169 proteins and 22,758 phosphosites with a localization probability >0.75. However, for further data analysis we only considered a stringently filtered dataset of 5,783 proteins and 7,387 phosphosites, respectively, with quantitative values in at least two out three biological replicates (“quantified”; Figure 1B, Supplementary table S1).

**Figure 1.**
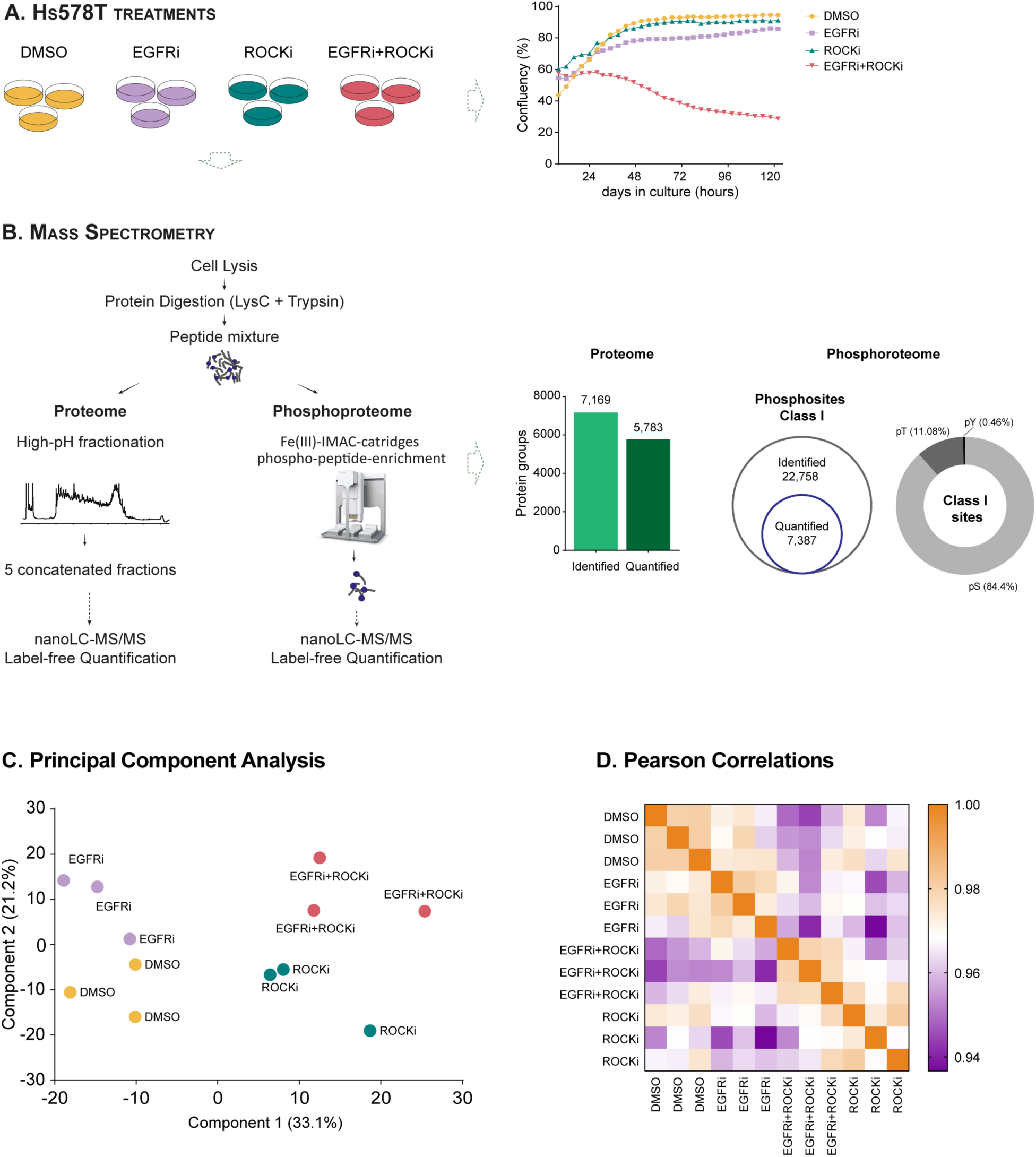
Experimental design and overview of the proteome data. **A.** Hs578T cells were treated respectively with DMSO, EGFRi (gefitinib), ROCKi (GSK269962A) or EGFRi+ROCKi combination treatments. Cell viability measured upon 120-h (5 days) treatments. **B.** Phospho(proteomics) workflow. After cell lysis, proteins were digested using LyC/trypsin. For in-depth proteome analysis, peptides were fractionated by high-pH reversed-phase chromatography and concatenated into 5 fractions prior to nanoLC-MS/MS analysis. Phosphopeptide enrichment was performed using an automated Fe(III)-IMAC workflow on the Bravo AssayMAP Platform. Bar plot of the total number of identified and quantified proteins. Total number of class I phosphosites identified and quantified. Distribution of Ser/Thr/Tyr phosphosites identified. **C.** Principal component analysis (PCA) and heatmap of the Pearson correlations coefficients based on their global proteomic expression profiles. Both PCA analysis and Pearson Correlation coefficients showed clustering of replicates and a clear separation between the different treatment conditions.

For an overall assessment of the effect of the four different treatment conditions on the global proteome profiles we employed principal component analysis (PCA). As can be seen in Figure 1C, all biological replicates of each condition clustered together, while principal component 1 (PC1) and principal component 2 (PC2) revealed a clear partition between the different treatments. In particular, PC1 clearly separated the DMSO and EGFRi treated samples from the ROCKi and EGFRi+ROCKi, while PC2 showed a segregation of the EGFRi and EGFRi+ROCKi from the rest and account for 33.1 % and 21.2% of the variability, respectively. The trend observed by PCA was confirmed by the Pearson correlation coefficients of the proteome data (Figure 1D). The Pearson correlations between the biological replicates and between the different treatments was >0.96 and >0.91, respectively.

### Comparative analysis between different treatment conditions revealed an additive effect of EGFR and ROCK inhibition upon combination treatment

Next, to address the statistical differences between the different drug treatments and obtain a view of potential functional proteomic changes, we performed an ANOVA test (FDR 5%), which identified 995 significantly changing proteins between any of the four conditions (Figure 2). Hierarchical clustering of these proteins revealed again a clear separation between the single inhibitor treatments (EGFRi and ROCKi) and a partial overlap with either of the two upon combination treatment of the Hs578T cells. In particular, the heatmap showed segregation of the ANOVA significant proteins into three main clusters; one, which was specific to proteins down-regulated in the EGFRi+ROCKi treatment (cluster A) and two that included up-regulated proteins that were common between the EGFRi+ROCKi treatment with the EGFRi (cluster B) and the ROCKi (cluster C) treatments, respectively (Figure 2). Gene ontology (GO) analysis of the proteins in cluster A revealed that EGFRi+ROCKi treatment resulted in down-regulation of nuclear and adhesion proteins, and subsequently down-regulation of biological processes including chromatin remodeling, mRNA processing and cell adhesion. Enrichment analysis of the proteins in cluster B showed that proteins up-regulated in the EGFRi and ROCKi treatments were involved in cellular processes including cholesterol biosynthesis, oxidation-reduction and autophagy, while proteins in cluster C (high expression in ROCKi and EGFRi+ROCKi) revealed strong enrichment of the terms cell adhesion, regulation of mRNA stability and intracellular protein transport. Interestingly, upon ROCKi and EGFRi+ROCKi treatments we observed an up-regulation of the proteasome complex, indicating the implication of another degradative pathway (together with autophagy) in the EGFRi+ROCKi treated cells.

**Figure 2.**
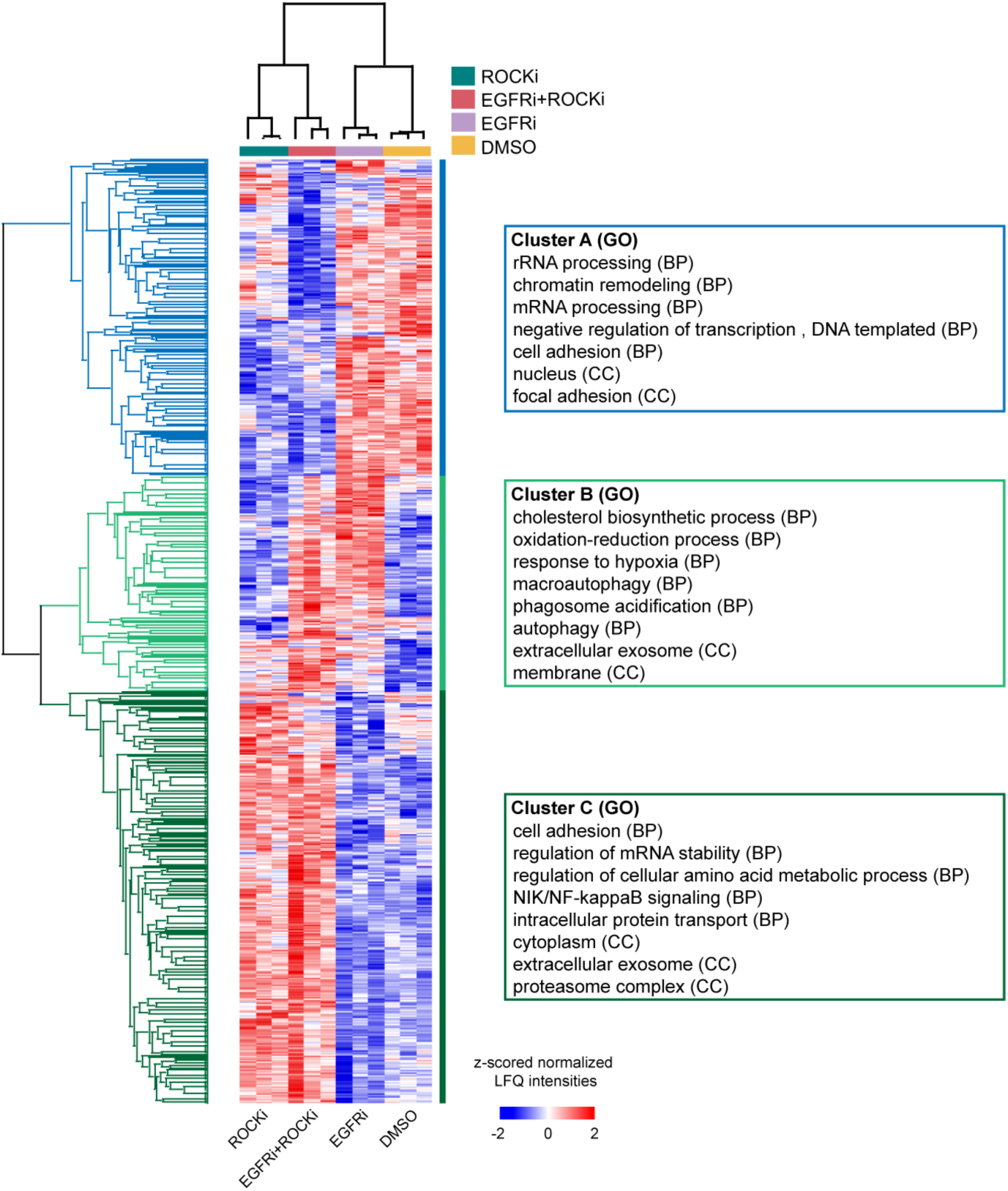
Proteins differentially expressed across different treatments. Heatmap showing relative protein expression values (z-scored and Log_2_-transformed LFQ protein intensities) of the differentially expressed proteins (ANOVA, FDR < 0.05) between the different samples after unsupervised hierarchical clustering. On the right, gene ontology analysis of proteins significantly down-regulated (Cluster A) and up-regulated (Clusters B and C) in EGFRi+ROCKi treatment.

In addition, comparative analysis performed on the phosphoproteome data revealed 1052 phosphosites to be differentially regulated (ANOVA test, p-value<0.05) upon the different treatments. In agreement to the proteome data, unsupervised hierarchical clustering of the regulated phosphosites and subsequently gene ontology (GO) analysis on the respective clusters showed a huge effect on processes related to transcription, cell-cell adhesion and cytoskeleton organization (both actin and microtubule cytoskeleton) upon EGFRi+ROCKi treatment (suppl. Fig. 1).

### Differentially expressed autophagy-related proteins in EGFRi, ROCKi and EGFRi+ROCKi treatments

The quantitative comparison between the different treatment conditions revealed a cluster of proteins involved in the process of autophagy to be significantly up-regulated upon single EGFR and dual EGFR and ROCK inhibition, indicating that autophagy might play a role in the TNBC cells’ response to therapy. Therefore, to further mine our quantitative data and provide insights into the molecular changes induced upon each treatment condition, we performed pairwise comparisons of each treatment condition (EGFRi-, ROCKi- and EGFRi+ROCKi-treated cells) to its untreated control (DMSO-treated cells) and focused on the differential regulation of autophagy-related proteins.

As shown in Figure 3, gefitinib treatment (EGFRi) compared to the DMSO-treated cells resulted in up-regulation of known autophagic markers. Amongst them, we detected the autophagy-related protein LC3 B (MAP1LC3B), which is used as a phagophore or autophagosome marker, and the GABARAP and GABARAPL2 proteins, which are involved in the later stages of autophagosome formation, in particular the phagophore elongation and closure [21]. Moreover, the cargo-specific autophagy receptors CALCOCO2, SQSTM1/p62 and its paralogue NBR1, the autophagy regulator TMEM59 and the lipid kinase PI4K2A, which plays a role in the autophagosome-lysosome fusion [22], were also up-regulated in response to gefitinib treatment.

**Figure 3.**
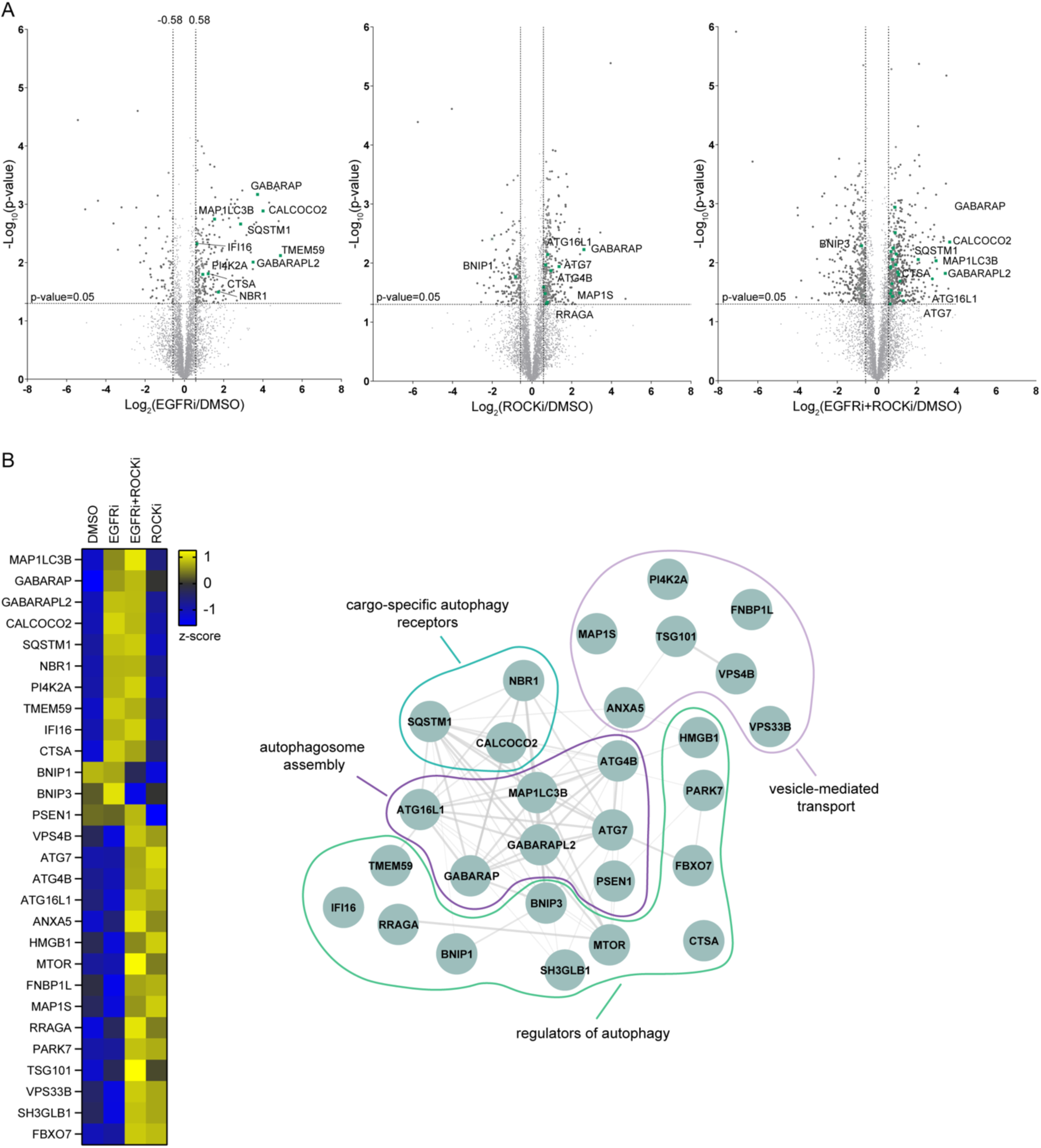
Autophagy-related proteins differentially expressed in EGFRi, ROCKi and EGFRi+ROCKi treatments. **A.** Volcano plots of the p-values vs the Log_2_ protein abundance differences between different treatment conditions; significantly enriched autophagy-related proteins are highlighted in green. **B.** Heatmap showing z-scored and Log_2_-transfromed LFQ protein intensities of the autophagy-related proteins across the different treatments and their protein-protein interaction network.

When we compared the proteomes of cells treated with ROCKi alone to the untreated control and looked for changes in the expression levels of known autophagy markers, we identified several proteins to be up-regulated upon ROCK inhibition. In particular, we identified the GABARAP receptor together with the autophagy-related (ATG) genes ATG7, ATG16L1 and ATG4B, which participate in the formation of phagophores and the initiation of autophagy [23]. Moreover, proteins with known roles in the regulation of autophagy such as the GTPase RRAGA, which activates autophagy in response to amino acids [24] and the microtubule-associated protein MAP1S, which is required for autophagosome trafficking along microtubular tracks [25] showed increased expression upon ROCKi treatment.

Finally, consistent with the ANOVA significantly changing proteins upon combination treatment, proteome expression changes in the EGFRi+ROCKi treated cells compared to the control cells (DMSO) revealed an accumulation of many autophagy-related proteins showing an enhanced and partly combinatorial effect of the regulation changes in the respective single treatments. An overview of the differentially regulated autophagy-associated proteins and their abundances across the different treatments is presented in the heatmap in Figure 3B, along with their protein-protein interaction network.

These findings further indicated the autophagy process as a potential pathway induced by the cells upon combination treatment, which might eventually cause cell death.

### EGFRi+ROCKi treatment induces autophagy in TNBC cells

To exclude that autophagy induction is a cell line-specific (Hs578T cells) response to the combinatorial treatment, and to determine whether it can be considered a general process in the response of TNBC cells, we decided to analyze the proteomic changes upon EGFRi+ROCKi treatment in another TNBC cell line. To this end, we chose the Cal51 cell line, whose sensitivity to EGFRi+ROCKi treatment has been previously reported as well [18]. Following the same experimental workflow as the Hs578T cells, the proteomes of Cal51 cells treated for 48h with DMSO, EGFRi, ROCKi and EGFRi+ROCKi were compared by label-free quantitation. Similar to the Hs578T cells, PCA analysis showed co-clustering of all the biological replicates of each treatment condition, except for one EGFRi-treated sample, which was an outlier and was therefore excluded from further analysis. In addition, the EGFRi- and DMSO-treated samples clustered close together, while ROCKi treated samples clustered with the EGFRi+ROCKi treated samples. These proteome similarities were also reflected in the Pearson correlations (Suppl. Fig. 2).

Furthermore, unsupervised hierarchical clustering of the 738 proteins, which were differentially expressed across the different treatment conditions (ANOVA, FDR<0.05) showed an enrichment in proteins involved in similar biological processes as those found on the Hs578T cells. For instance, treatment of Cal51 cells with ROCKi and EGFRi+ROCKi resulted in down-regulation of nuclear proteins involved in transcription, rRNA processing and cell cycle. Additionally, treatment of Cal51 cells with single EGFR and combined EGFR+ROCK inhibitors showed up-regulation of proteins involved in autophagy, while ROCKi and EGFRi+ROCKi treatments resulted in up-regulation of cell-cell adhesion, protein transport and an increase in the expression levels of protein of the proteasome complex (Suppl. Fig. 3A). Moreover, the specific autophagy-related proteins and their differential regulation across each treatment condition are highlighted in the respective volcano plots (Suppl. Fig. 3B). Again, an additive effect on the expression of autophagy related proteins is observed in the proteome changes induced upon combination treatment.

We next set out to validate our mass spectrometry data and the induction of autophagy by western blot analyses in a panel of TNBC cell lines. Following the induction of autophagy, the microtubule-associated protein LC3, MAP1LC3 (MAP1LC3-I) is converted to membrane bound phosphatidylethanolamine (PE)-conjugated MAP1LC3 (MAP1LC3-II), and the expression of MAP1LC3-II is frequently used as a phagophore or autophagosome marker [26]. We therefore evaluated the expression levels of this protein as well as those of p62 (SQSTM1), which as mentioned previously is a known autophagy receptor that links ubiquitinated proteins to MAP1LC3. As shown in Figure 4A, in MDAMB231 cells, EGFR inhibition compared to DMSO treatment resulted in an increase in the protein levels of the MAP1LC3-II and the p62 (SQSTM1) proteins, indicating autophagy induction. When cells were treated with EGFRi+ROCKi the protein levels of LC3-II and p62 remained high, whereas ROCKi treatment alone did not have any influence on autophagy. Interestingly, although combined inhibition of EGFR and ROCK led to increased AMPK phosphorylation, which is another indication of autophagy induction, p-AMPK was not detected in cells treated with EGFRi alone. Finally, in MDA-MB-231 cells, we observed a steep decrease in phosphorylated rpS6 levels upon ROCK inhibition while, rpS6 phosphorylation declined even further in the case of combined EGFRi+ROCKi treatment, which is indicative of an inactive state of the mTOR pathway [27] and which has been previously associated with cell growth inhibition and cell cycle arrest induction [28]. The drug-induced changes in the levels of LC3-II protein and the phosphorylation status of rpS6 were reproducible in the triple-negative cell lines Hs578T, Cal120 and HCC1806 (Figure 4B).

**Figure 4.**
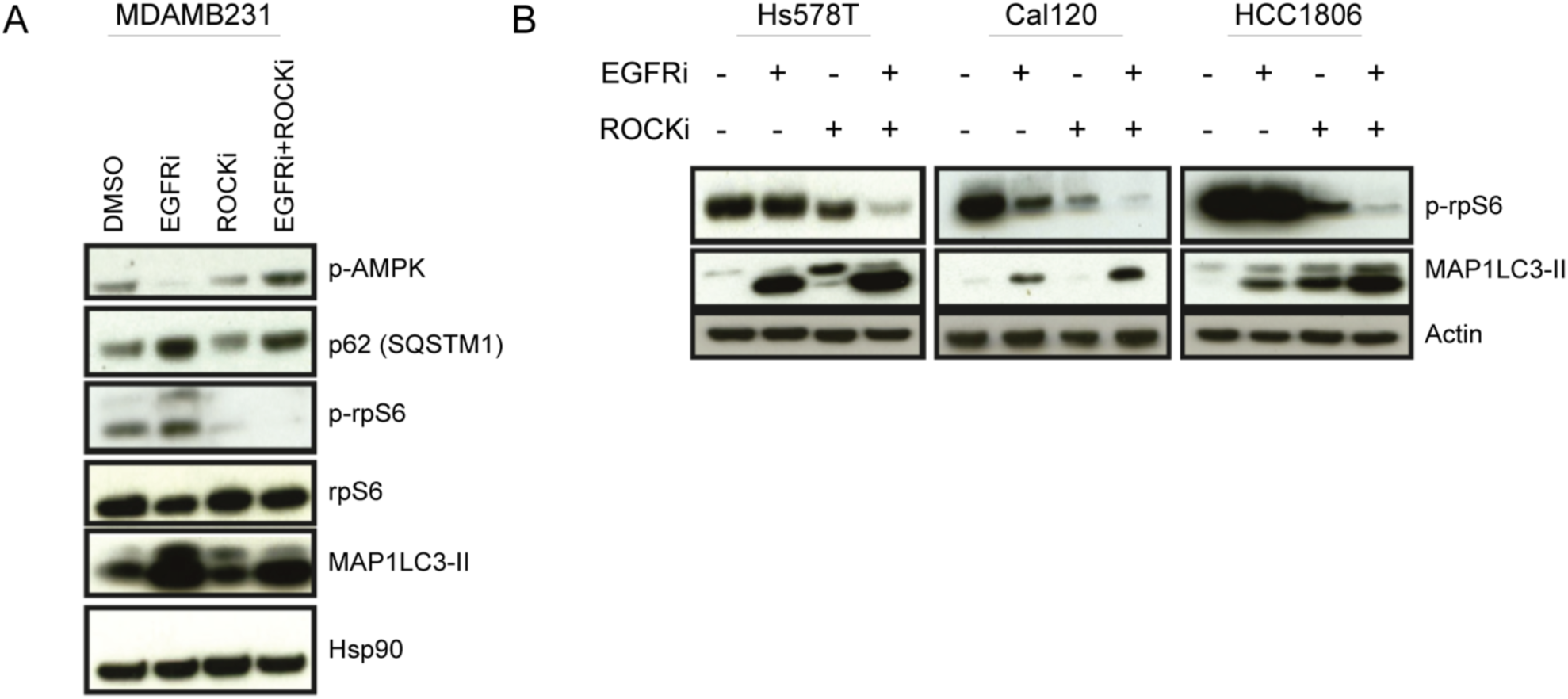
Autophagy induction in TNBC cells. **A.** Western blot analyses to evaluate the protein expression levels of known autophagy markers and rpS6 phosphorylation in DMSO, EGFRi, ROCKi and EGFRi+ROCKi treated MDA-MB-231 cells. Hsp90 is used as a loading control. **B.** Western blot analysis of LC3-II (MAP1LC3-II) and phosphorylated rpS6 levels in Hs578T, Cal120 and HCC1806 cells. Actin is used as a loading control.

### EGFRi+ROCKi treatment impairs autophagic flux

The results presented thus far provide compelling evidence that combined inhibition of EGFR and ROCK triggers autophagy induction in triple-negative breast cancer cells, however, how autophagy induction leads to cell death remains unclear. Therefore, in a next step we decided to monitor autophagic activity in live cells treated with DMSO, EGFRi, ROCKi and EGFRi+ROCKi, respectively.

To do so, treated Hs578T cells were stained with the Cyto-ID autophagy green dye, which specifically labels autophagic vacuoles, and were visualized using live cell imaging in the IncuCyte System (Suppl. Fig. 4). The dye enables clear detection and quantification of autophagic and pre-autophagic vacuoles that directly correlate with induction of autophagy [29]. The control group (DMSO) exhibited faint Cyto-ID green fluorescence while EGFR inhibition by gefitinib treatment induced the appearance of green autophagic vacuoles in the cells. By contrast, no significant autophagy was identified in the ROCKi-treated cells. Moreover, in the combination treatment, inhibition of ROCK activity did not abolish the autophagy induction mediated by gefitinib treatment, but instead resulted in an increased number of stained autophagic vacuoles compared to the EGFRi-treated cells. This accumulation of autophagic vacuoles in the EGFRi+ROCKi-treated cells can either be a result of increased stimulation of autophagy resulting in rapid formation of autophagic vacuoles or due to inefficient autophagosome turnover and clearance caused by impaired autophagosome-lysosome fusion. Thus, in order to distinguish between these two possibilities, we assessed the autophagic flux by monitoring the accumulation of autophagic compartments induced by chloroquine (CLQ), a lysosome inhibitor that blocks the fusion of autophagosomes and lysosomes [30]. As shown in Figure 5, combined treatment of EGFRi+CLQ resulted in a significant increase in the number of autophagic vacuoles compared to EGFRi alone, due to impaired autophagic flux. The addition of CLQ to EGFRi treatment raised the number of observed autophagic vacuoles to a similar level as observed for our EGFRi+ROCKi combination treatment. On the other hand, co-administration of EGFRi+ROCKi with CLQ did not cause a significant increase in the autophagic vacuoles formation compared with the EGFRi+ROCKi treatment alone. These results indicate that EGFRi+ROCKi does not stimulate autophagic flux in Hs578T cells, beyond to that seen by single inhibition, and that the increase in autophagic vacuoles is caused by impaired autophagosome clearance instead of increased vacuole formation.

**Figure 5.**
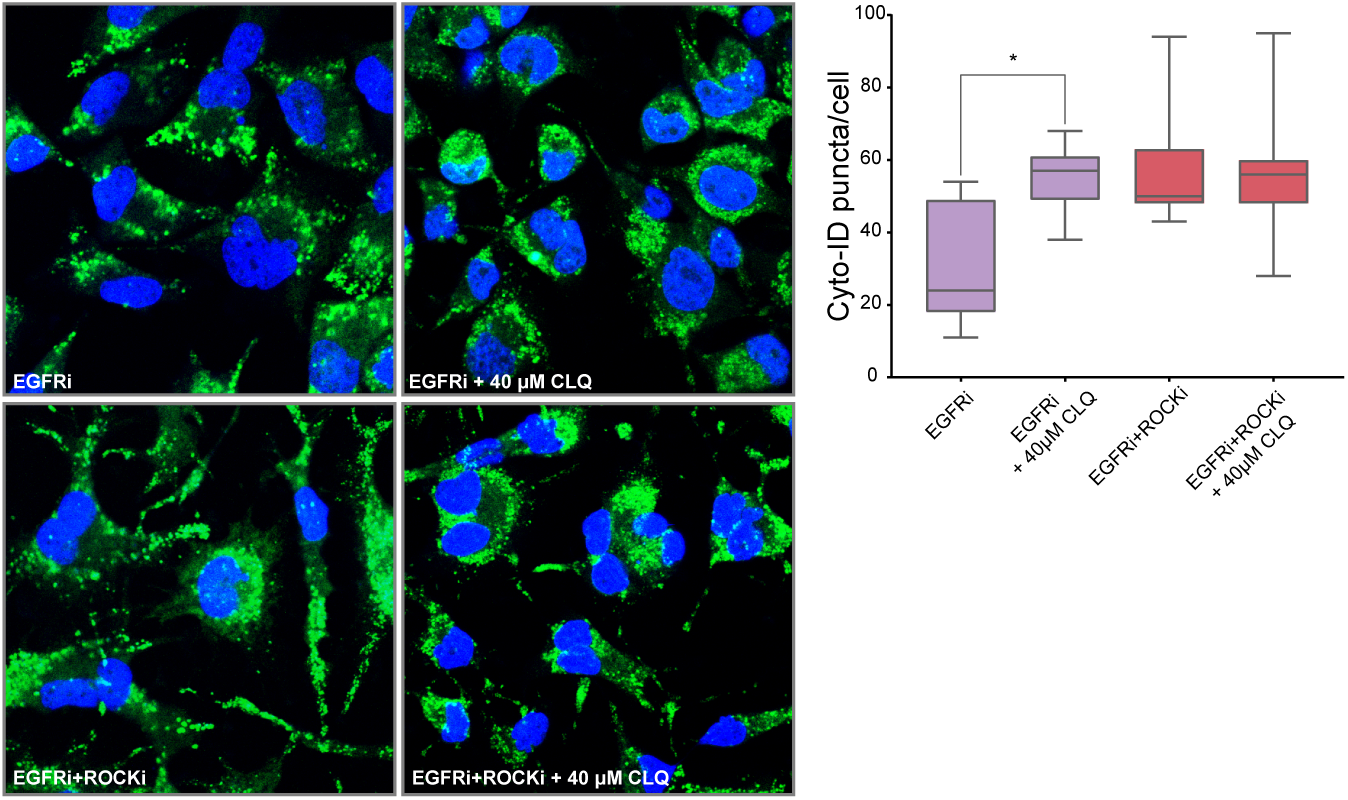
Autophagic flux in EGFRi and EGFRi+ROCKi-treated cells. EGFRi- and EGFRi+ROCKi-treated Hs578T cells, were incubated for 2 hours in the absence or presence of 40μM chloroquine (CLQ) and subsequently stained with the Cyto-ID dye (green). Nuclei were counter-stained with Hoechst 33342 dye (blue). Images were obtained by confocal microscopy and autophagic vacuoles were counted to assess autophagic flux per treatment condition. The graph on the right shows the average of Cyto-ID puncta per cell (n= 15,*p-value <0.01 from two-sided, unpaired *t*-test).

Taken together, these findings strongly indicate that cell death upon EGFRi+ROCKi treatment occurred due to autophagy blockade and impairment of the EGFRi-induced autophagic flux. Thus, ROCK activity is essential for efficient autophagy process.

### ROCK-associated cytoskeletal changes affect autophagy

As inhibition of ROCK activity led to impaired autophagy, we next set out to investigate the mechanisms underlying this effect. As revealed by the gene ontology enrichment analysis of the ANOVA significant proteins and phosphosites (Figure 2 and Suppl. Fig. 1), ROCK inhibition had a substantial effect on the expression levels of several cytoskeletal and focal adhesion proteins. This finding is consistent with the involvement of ROCK in the regulation of cell shape and movement [31] and was also evident from the major morphological changes that occurred in the cells upon ROCKi and EGFRi+ROCKi treatment, where cells became flattened and acquired neuron-like long extensions. Recent evidence indicates an important role of actin cytoskeleton dynamics together with myosin motor proteins in the various steps of the autophagy process, ranging from the early stages of phagophore formation and expansion, to autophagosome trafficking and fusion with the lysosome [32, 33]. In line with these findings, we observed marked changes in the expression levels of proteins involved in focal adhesion and the regulation of the actin and microtubule cytoskeleton, which were down- and up-regulated, respectively (Figure 6), upon combination treatment.

**Figure 6.**
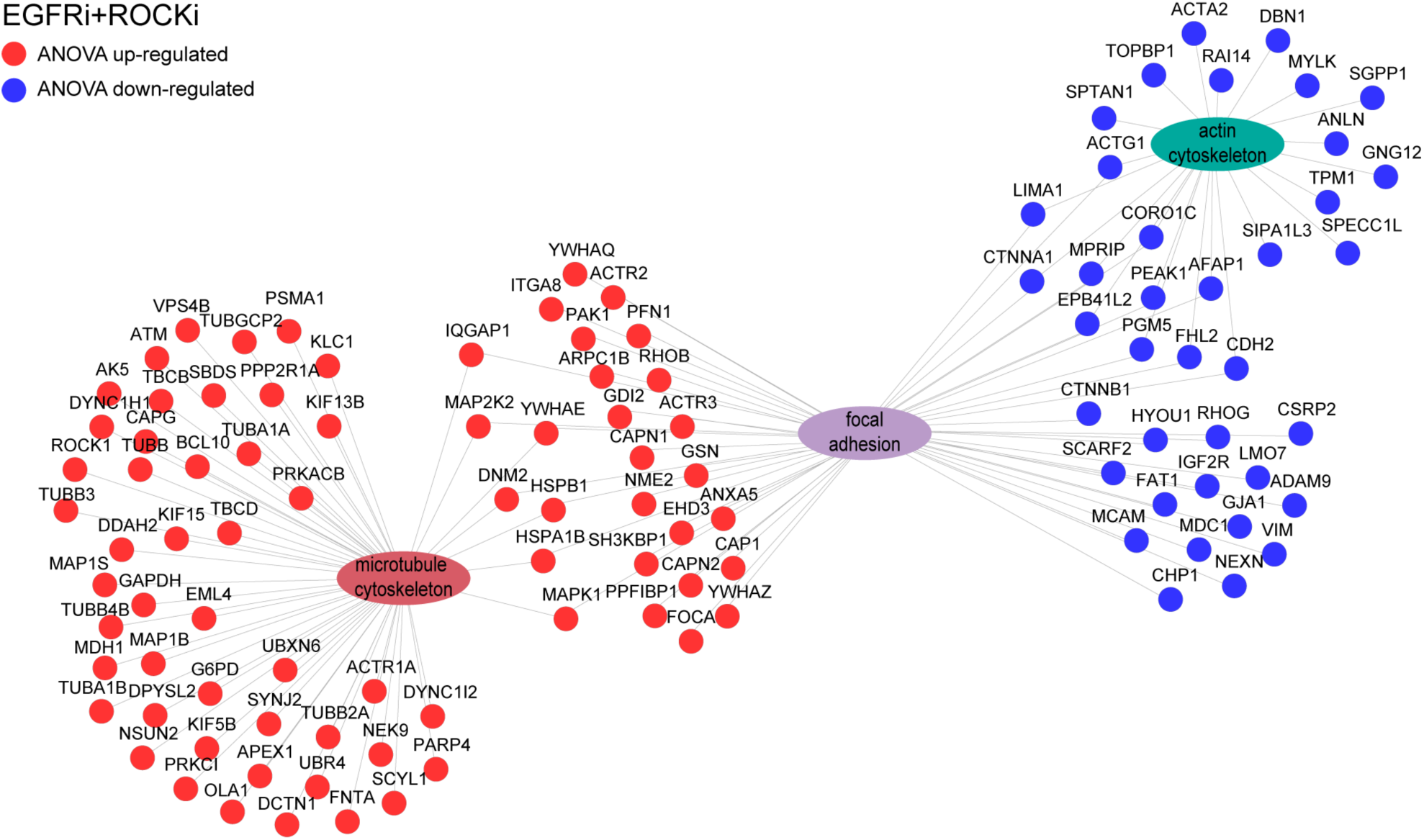
Overview of the differentially expressed cytoskeleton-related proteins upon ROCK activity inhibition. Functional enrichment network of the ANOVA significant up- (red) and down-regulated (blue) cytoskeleton-related proteins in the combination treatment using the *ToppCluster* tool (FDR correction, p-value < 0.05) [48].

Actin filament networks have been previously suggested to have a scaffolding role in generating the shape of the phagophore with the recruitment of the Arp2/3 complex that is known to promote actin branching and polymerization inside the expanding phagophore [34-36]. Interestingly, in our data we observed up-regulation of the actin related proteins ACTR2, ACTR3 and ARPC1B, in the EGFRi+ROCKi treated cells, which are core subunits of the Arp2/3 complex. On the other hand, the actins, alpha-actin-2 (ACTA2) and gamma-actin (ACTG1), together with several actin binding proteins such as the tropomyosin alpha-1 chain (TPM1), the drebrin (DBN1), the actin filament associated protein 1 (AFAP), the coronin-1C (CORO1C), nexilin (NEXN), which are essential in stabilizing cytoskeleton actin filaments, were down-regulated. Furthermore, amongst the down-regulated proteins upon EGFRi+ROCKi treatment, we detected proteins involved in actomyosin-based motility such as the proteins anillin (ANLN), the myosin light chain kinase (MYLK) and the myosin phosphatase Rho interacting protein (MPRIP) that regulate actin-myosin interactions as well as proteins playing a role in actin cytoskeleton and microtubule stabilization such as cytospin-A (SPECC1L) and the formin-binding protein 1-like (FNBP1L).

It has been shown that autophagosome movement in the cytoplasm is dependent on microtubules [37]. Once the autophagosomes are formed, they move along microtubular tracks towards the microtubule-organizing center where lysosomes are enriched [38]. Here, upon combination treatment, we found several tubulins and tubulin-associated proteins to be up-regulated, including the microtubule-associated proteins MAP1S and MAP1B, which interact with LC3-I and LC3-II and recruit them to stable microtubules [25]. However, in contrast to the proteome data, our phosphoproteome data showed a significant down-regulation on the phosphorylation status of several proteins involved in the microtubule cytoskeleton organization. Within the identified down-regulated phosphosites, we found multiple sites of the microtubule associated proteins MAP1A (S2022), MAP1S (T638, S638), MAP1B (S2098, S832, S831, S1144, T1932) and MAP4 (S941, S507, S510). Although the exact functionality of the different phosphosites has not been yet investigated, several lines of evidence suggest that the binding of MAPs to microtubules is regulated by phosphorylation [39].

### Crosstalk of autophagy and UPS

Interestingly, upon inhibition of autophagy by ROCKi we observed a significant up-regulation on the expression levels of core and regulatory subunits of the proteasome, implicating a potential activation of the ubiquitin-proteasome system (UPS) as a compensatory mechanism of cells to reduce the burden of accumulated autophagic substrates. A protein network of the detected multiple proteasomal subunit proteins (ANOVA significant) and their increased abundance levels in ROCKi- and EGFRi+ROCKi-treated cells compared to the DMSO and EGFRi treatments are illustrated in Figure 7A. Amongst them, we detected proteins of the *α*- and *β*- subunits of the 20S core structure of the proteasome in the mammalian cells, including the catalytic proteasome β1 subunit, PSMB1. However, although our proteome data indicate a potential crosstalk between the two major degradation systems, complementary experiments to measure whether proteasomal activity is indeed enhanced upon EGFRi+ROCKi treatment need to be performed to confirm our hypothesis.

**Figure 7.**
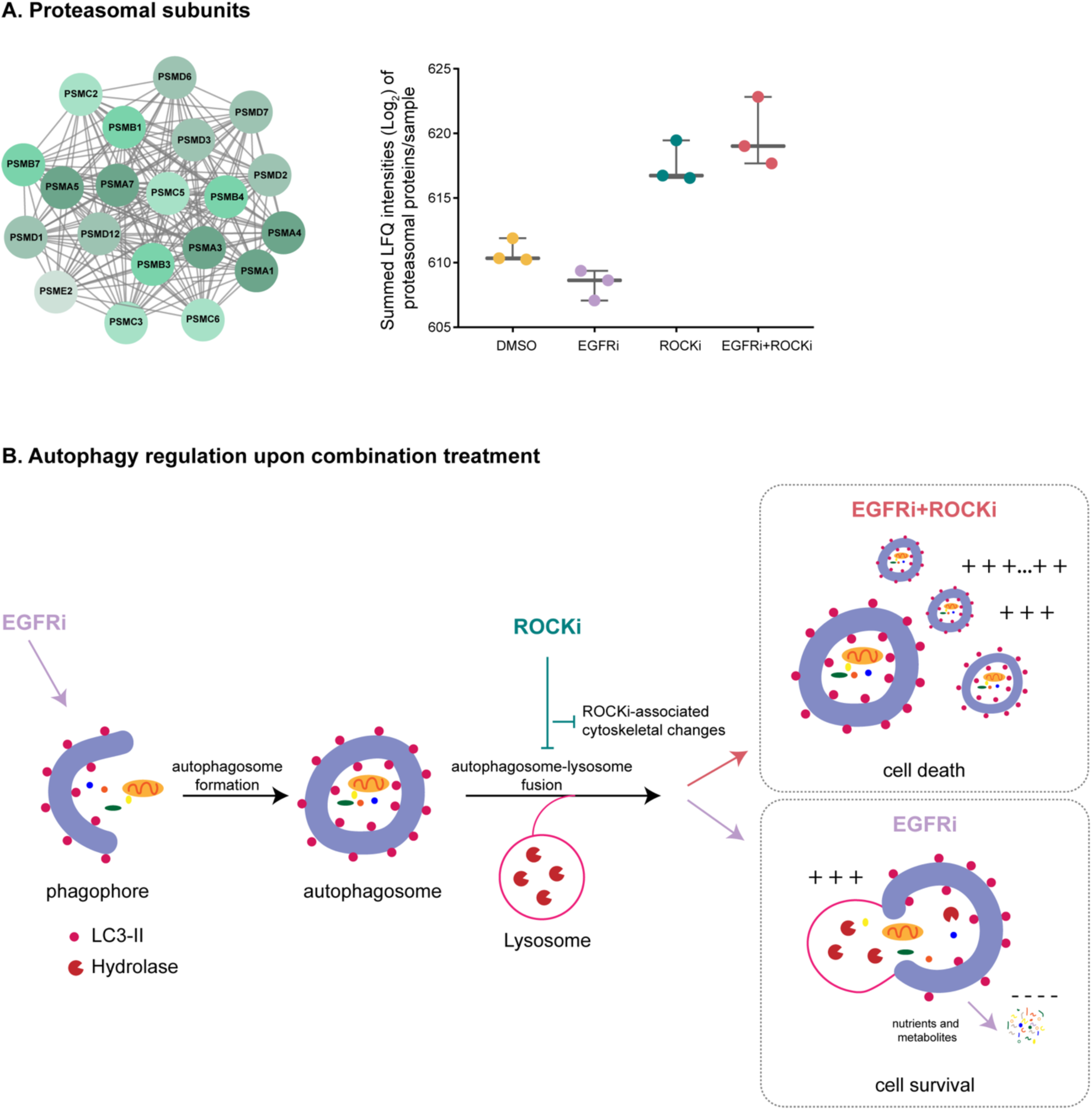
Regulation of proteasomal subunits and autophagy upon EGFR and/or ROCK inhibition. **A.** String network of all the identified proteasomal subunit proteins and their relative abundance across the different treatments displayed by summing their Log_2_-transformed LFQ intensities. **B.** EGFR inhibition induces autophagic flux in TNBC cells, which can be impaired upon co-inhibition of ROCK activity leading to an increased accumulation of autophagic vacuoles and possibly autophagic cell death.

## Discussion

Triple-negative breast cancer is an aggressive BC subtype, which suffers from the absence of known drug targets and is associated with an especially poor prognosis [40]. Based on previous work [18], which revealed that the combination of EGFR and ROCK inhibitors effectively reduced TNBC cell growth by inducing cell cycle arrest, we here complement these previous findings by providing an insight into the molecular mechanisms triggered by the combinatorial EGFRi+ROCKi treatment.

Using a quantitative (phospho)proteomics approach to compare the proteome changes upon single and combination treatments, we identify autophagy activation as a potential mechanism of the cells’ response to treatment. We show that EGFR inhibition by gefitinib induces autophagy activation in TNBC cells, which was evident from the increased expression levels of several autophagy protein markers (e.g. MAP1LC3, GABARAP) and the formation of autophagic vacuoles. Moreover, we found that co-inhibition of EGFR and ROCK causes accumulation of autophagic vacuoles in triple-negative breast cancer cells and subsequent findings revealed that this accumulation is caused by the inhibition of autophagic flux as a result of ROCK activity inhibition. EGFR inhibitors have previously been associated with autophagy regulation, although the specific function of this induction in cancer remains biphasic. In some studies, autophagy induction serves a cytoprotective response in cancer cells, while other studies report that enhanced autophagy after treatment of EGFR inhibitors can result in autophagic cell death [41]. In this study, our results suggest that autophagy induction has a pro-survival role in triple-negative breast cancer cells upon gefitinib treatment.

In addition to the contradicting literature regarding EGFR inhibitors and autophagy, ROCK activity has also been linked to this process, albeit with conflicting opinions regarding its function. Inhibition of ROCK activity can lead to autophagy impairment and cell death [42], while ROCK activity is required for starvation-mediated autophagy, since ROCK inhibition resulted in a decreased number of autophagosomes in cells under starvation conditions [43]. Conversely, a study by Mleczak *et a*l., showed that ROCK activity inhibited autophagy while the opposite was true for ROCK inhibition, which enhanced the autophagy response upon starvation and led to the accumulation of enlarged early autophagosomes that matured into enlarged late degradative autolysosomes [44]. Here, we show that ROCK activity is required for gefitinib-induced autophagy and that inhibition of ROCK leads to autophagy blockade and accumulation of autophagic vacuoles due to impaired autophagosome clearance. Given the key roles of the cytoskeleton in the different stages of autophagy, we speculate that the ROCKi-associated cytoskeletal changes are responsible for the blockage of autophagy. Indeed, our proteome data revealed major expression changes associated to the actin- and microtubule- cytoskeleton, which could cause a block during various steps of the autophagic pathway from the early stages of phagophore formation and expansion to vesicle trafficking and fusion with the lysosomes. In line with this reasoning, we found that the number of observed autophagic vacuoles in our EGFRi+ROCKi combination treatment in the TNBC cells was at a similar level as after the addition of CLQ, which blocks the fusion of autophagosomes and lysosomes, to EGFRi single treatment.

Interestingly, upon ROCK inhibition, and subsequently autophagy impairment, we observed a significant up-regulation of several proteasomal subunit proteins indicating a potential link between the autophagy and the ubiquitin-proteasome system, which is the other major intracellular pathway for protein degradation in mammalian cells. Indeed, in agreement with our findings, extensive evidence indicates that connections and crosstalk exist between the two systems, which are interconnected and inhibition of one system leads to a compensatory up-regulation of the other system [45-47]. However, although our proteome data indicates up-regulation of the proteasome upon autophagy inhibition, as discussed above, further experiments to assess the increased proteasomal activity *in vitro* are necessary to confirm the crosstalk.

In summary, this work provides proteomic and functional evidence for the activation of autophagy as a survival mechanism to EGFR inhibition in triple-negative breast cancer cells, which can be impaired upon co-inhibition of ROCK activity and ultimately lead to TNBC cell death (Figure 7B). We therefore, believe that our data support the clinical potential of therapeutically inhibiting autophagy for improved cancer therapy.

## Acknowledgements

This work was supported by the Netherlands Organization for Scientific Research (NWO) through a VIDI grant for M.A. (723.012.102) and Proteins@Work, a program of the National Roadmap Large-scale Research Facilities of the Netherlands (project number 184.032.201) and the Dutch Cancer Society (10304) for S.R., M.A. and D.S.P.

## Author Contributions

Conceptualization, D.S.P. and M.A.; Methodology, S.R., S.I., S.v.D. and M.A.; Investigation, S.R., and M.A.; Writing – Original Draft, S.R. and M.A.; Writing – Review & Editing, S.R, D.S.P. and M.A.; Funding Acquisition, D.S.P. and M.A.

## Materials and methods

### Cell culture and inhibitors

MDA-MB-231 and Cal120 cells were cultured in Dulbecco’s modified Eagle’s medium (DMEM) supplemented with 10% Fetal Bovine Serum (Sigma), 2mM glutamine, 0.1mg/ml penicillin and 0.1ml/ml streptomycin (Gibco). HCC1806 and Hs578T cells were maintained in RPMI supplemented with glutamine. All cells were maintained in a humidified incubator at 37°C and 5% CO_2_. All cell lines were obtained from ATCC and have been regularly tested for mycoplasma contamination. For the (phospho)proteomics and western blot experiments, drugs were added on the following day of seeding. Cells were treated with the inhibitors Gefitinib (EGFRi, MedChem) or GSK269962A (ROCKi, Axon) or their combination (EGFRi+ROCKi) using the following concentrations: Hs578T, Cal51, MDA-MB-231, Cal120 and HCC1806 cells were treated with 20μM EGFRi. ROCKi concentrations were the following: for Hs578T 1.2μM, for Cal51 12μM, for MDA MB-231 4.8μM, for HCC1806 2.4μM and for Cal120 was 30μM.

### Sample preparation for mass spectrometry

Cal51 and Hs578T cells were harvested in triplicates in cold PBS after a 2-day treatment with DMSO, EGFRi, ROCKi or combination (EGFRi+ROCKi). The cellular pellets were resuspended in lysis buffer containing 1% (w/v) sodium deoxycholate (SDC), 10 mM TCEP, 40 mM chloroacetamide, 100 mM Tris, pH 8.5, supplemented with 1 tablet of Complete mini EDTA-free mixture (Roche) and 1 tablet of PhosSTOP phosphatase inhibitor cocktail mixture (Roche) per 10ml of lysis buffer, and subsequently lysed by boiling for 5 min at 95°Cand sonication (Bioruptor, model ACD-200, Diagenode) for 15 min at level 5 (30 sec ON, 30 sec OFF). Cell debris was then removed by centrifugation at 20,000 × g for 15min at 4°C. Prior to in-solution digestion, the total protein concentration was quantified by Bradford assay (Bio-Rad). For label-free quantification, input amounts were normalized based on the total protein contents (50 μg of total protein lysate per sample). The lysate was diluted 1:10 with 50 mM ammonium bicarbonate for Lys-C and trypsin digestion. Protein digestion was performed overnight at 37 °Cwith Lys-C (Wako) at an enzyme/protein ratio 1:75 and trypsin (Sigma) at an enzyme/protein ration of 1:50. The digest was acidified by adding 4% formic acid (FA) to precipitate SDC and samples were subsequently desalted using Sep-Pak C18 cartridges (Waters Corporation) and further submitted to phosphorylation enrichment or high pH fractionation for in-depth proteome analysis.

### High-pH reversed-phase fractionation

50 μg of peptides of each sample were reconstituted in 10mM ammonium hydroxide, pH 10 and loaded on a Gemini 3µm C18 110 Å 100 × 1.0 mm column (Phenomenex) using an Agilent 1100 binary pump (Agilent Technologies). The peptides where concentrated on the column at 100 µl/min using 100% buffer A (10mM Ammonium Hydroxide, pH 10) for 2 minutes after which the fractionation gradient initiated as follow: 5% solvent B (10mM ammonium Hydroxide in 90% ACN, pH 10) to 30% B in 53 minutes, 70% B in 7 minutes and increased to 100% B in 3 minutes at a flow rate of 100 µl/min. In total 60 fractions of 1 minute were collected using an Agilent 1260 infinity fraction collector, and were pooled into 5 fractions using the concatenation strategy as described [19]. The pooled fractions were dried in a vacuum centrifuge and stored at −80°Cuntil further analysis.

### Mass Spectrometry analysis

Nanoflow LC-MS/MS data were acquired on an Orbitrap Q Exactive Plus (Thermo Fisher) coupled to an Agilent 1290 Infinity UHPLC (Ultra-High Pressure Liquid Chromatography) system (Agilent Technologies). Both the trap (Dr Maisch Reprosil C18, 3 μm, 2 cm x 100 μm) and the analytical (Agilent Poroshell EC-C18, 2.7 μm, 50 cm x 75 μm) columns were packed in-house. Peptides were trapped for 10 min at 5 μl/min in 100% solvent A (0.1 M acetic acid in water). Separation was performed at a column flow rate of ∼300 nl/min (split flow from 0.2 ml/min) and the gradient was as follows: 13% up to 40% solvent B (0.1 M acetic acid in 80% acetonitrile) in 95 min, 40-100% in 3 min and finally 100% for 1 min. The mass spectrometer was programmed in the data-dependent acquisition mode. Full scan MS spectra from *m/z* 375-1,600 were acquired at a resolution of 35,000 with an automatic gain control (AGC) target value of 3e6. The 10 most intense precursor ions were selected for fragmentation using HCD. MS/MS spectra were obtained at a 17,500 resolution with an AGC target of 5e4. HCD fragmentation was performed at a normalized collision energy (NCE) of 25%.

### Phosphopeptide enrichment and MS analysis

Phosphopeptide enrichment was performed using a combination of Fe(III)-IMAC cartridges and an automated setup, the AssayMAP Bravo Platform (Agilent Technologies) as described previously [20]. Briefly, Fe(III)-NTA cartridges were primed with 250 μL of 0.1% TFA in ACN and equilibrated with 250 μL of loading buffer (80% ACN/0.1% TFA). Peptides were dissolved in 200 μL of loading buffer and loaded onto the cartridge. The columns were washed with 250 μL of loading buffer, and the phosphorylated peptides were eluted with 25 μL of 1% ammonia directly into 25 μL of 10% formic acid. Subsequently, the samples were dried down in a vacuum centrifuge. Next, phosphopeptides were reconstituted in loading buffer containing 10% formic acid and analyzed by nanoLC-MS/MS on a Q Exactive HF (ThermoFisher Scientific) coupled to an Agilent 1290 Infinity System (Agilent Technologies). As previously described, eluted phosphopeptides were delivered to a trap column (100 μm i.d. × 2 cm, packed with 3 μm C18 resin, Reprosil PUR AQ, Dr. Maisch) at a flow rate of 5 μL/minute in 100% loading solvent A (0.1% FA, in HPLC grade water). After 10 min of loading and washing, peptides were transferred to an analytical column (75 µm i.d. x 50 cm, packed with 2.7 µm Poroshell 120 EC C18, Agilent Technologies) and eluted at room temperature using an 95 min with an LC gradient from 8% to 32% solvent B (0.1% FA, 80% ACN). The Q Exactive HF was operated in data-dependent acquisition mode using the following settings: full-scan automatic gain control (AGC) target 3e6 at 60,000 resolution; scan range 375–1600 m/z; Orbitrap full-scan maximum injection time 20 ms; MS2 scan AGC target 1e5 at 30,000 resolution; maximum injection time 50 ms; normalized collision energy 27; dynamic exclusion time 16s; isolation window 1.4 m/z; 12 MS2 scans per full scan.

### Data Processing

Raw MS files were processed with MaxQuant (version 1.6.2.3) [49]. The Andromeda search engine [50] was used to search the MS/MS data against the forward and reverse Human Uniprot database (20,386 entries, version August 2018). Trypsin/P was specified as enzyme allowing up to two missed cleavages. Cysteine carbamidomethylation was set as a fixed modification, while methionine oxidation and protein N-term acetylation were set as variable modifications. For the phosphoproteome analysis, serine, threonine and tyrosine were selected as variable modification. Peptide spectrum match (PSM) and protein identifications were filtered using a target-decoy approach at false discovery rate (FDR) of 1%. Label free quantification (LFQ) was performed using the MaxLFQ algorithm [51] integrated into MaxQuant with the following parameters: LFQ minimum ratio count was set to 2, the Fast LFQ option was enabled, LFQ minimum number of neighbors was set to 3, and the LFQ average number of neighbors to 6. The “match between runs” feature was enabled with a match time window of 0.7 min and an alignment time window of 20 min.

### Data analysis

All data were analyzed using the Perseus software [52]. LFQ intensities extracted by MaxQuant were Log2 transformed. The samples were grouped in triplicates (DMSO, EGFRi, ROCKi and EGFRi+ROCKi) and identifications were subsequently filtered for proteins having at least 2 valid values in at least one treatment group. Missing values were imputed on a basis of normal distribution with a downshift of 1.8 SDs and a width of 0.3 SDs, enabling statistical analysis. Only class I phosphorylation sites (localization probability p > 0.75) were used in subsequent phosphoproteome analyses. For hierarchical clustering, logarithmized LFQ intensities were first z-scored and subsequently clustered using Euclidean as a distance for column and row clustering. Principal component analysis (PCA) was performed using Perseus’ built-in tool. Differences in the protein levels between the different treated samples were calculated using an ANOVA test followed by a Benjamini-Hochberg multiple testing correction with a 5% FDR. Gene ontology (GO) analyses were performed with Database for Annotation, Visualization and Integrated Discovery (DAVID) v6.8 [53, 54]. Protein-protein interaction network analysis was performed using the Cytoscape StringApp [55, 56].

### Western blot analysis

Cells were harvested in ice by scraping in ice cold 1X PBS and the pellets were lysed in RIPA buffer (50 mM TRIS pH 8.0, 150 mM NaCl, 1% Nonidet P40, 0.5% sodium deoxycholate, 0.1% SDS, complete protease inhibitor cocktail (Roche), and phosphatase inhibitors 10 mM NaF, 1 mM Na_3_VO_4_, 1 mM sodium pyrophosphate, 10 mM beta-glycerophosphate). After sonication and centrifugation the protein concentrations were determined using the Bio-Rad protein assay (Bio-Rad). Equal protein amounts were loaded on 4-12% Bis-Tris polyacrylamide-SDS gels (NuPAGE) and transferred on to nitrocellulose membranes (Amersham). Membranes were blocked in 4% skimmed milk powder dissolved in 0.2% Tween-containing 1X PBS and incubated with primary antibodies followed by secondary antibodies (Invitrogen). Primary antibodies used were LC3 (5F10, Nanotools), p62 (610832, BD Biosciences), AMPK_Thr172_ (40H9, Cell Signaling), rpS6 (5G10, Cell Signaling), rpS6_Ser235/236_ (Cell Signaling), Hsp90 (sc-7947, Santa Cruz), Actin (AC-74, Sigma).

### Autophagy detection and quantification by Cyto-ID staining

Autophagy was measured using the CYTO-ID Autophagy Detection Kit (Enzo Life Sciences), according to the manufacturer’s detailed instructions provided. Briefly, 6×103 Hs578T cells were seeded in 96-well plates overnight and then treated with the respective drug treatments (DMSO, EGFRi, ROCKi, EGFRi+ROCKi) on the following day. After 20 h of treatment, cells were stained with the Cyto-ID dye and the green fluorescent autophagic vacuoles were subsequently visualized using the IncuCyte System (Essen Bioscience). Acquired images were analyzed using the IncuCyte software and green fluorescent objects were counted enabling the “Top-Hat” feature, which estimates and subtracts local background from the image. To monitor autophagic flux, cells were analyzed by confocal microscopy. Approximately, 25×103 cells were seeded on glass bottom imagining dishes (μ-Dish 8 well, Ibidi) and after overnight incubation with the single and combination treatments, addition of 40 μM chloroquine (Enzo Life Sciences) followed. Subsequently, cells were stained with the Cyto-ID Green Detection Reagent and the Hoechst 33342 Nuclear Stain (Enzo Life Sciences), fixed for 15 min with 4% paraformaldehyde (PFA) and subsequently analyzed by a confocal laser scanning microscope Carl Zeiss LSM 700 with a 40x oil-immersion objective lens. Green puncta in confocal images were quantified by ImageJ in combination with the ComDet (pluginhttps://github.com/ekatrukha/ComDet/wiki). Significant differences between the CLQ treated and untreated samples were analyzed using student’s t-test (two-tailed) to compare the two groups, with a p-value <0.05 to be considered significant.

### Data availability

All mass spectrometry proteomics data have been deposited to the ProteomeXchange Consortium via the PRIDE partner repository with the dataset identifier PXD013821.

